# *In vivo* photopharmacology with a caged mu opioid receptor agonist drives rapid changes in behavior

**DOI:** 10.1101/2021.09.13.460181

**Authors:** Xiang Ma, Desiree A. Johnson, Xinyi Jenny He, Aryanna E. Layden, Shannan P. McClain, Jean C. Yung, Matthew R. Banghart

## Abstract

Photoactivatable drugs and peptides can drive quantitative studies into receptor signaling with high spatiotemporal precision, yet few are compatible with behavioral studies in mammals. We developed CNV-Y-DAMGO, a caged derivative of the mu opioid receptor-selective peptide agonist DAMGO. Photoactivation in the mouse ventral tegmental area produced an opioid-dependent increase in locomotion within seconds of illumination. These results demonstrate the power of *in vivo* photopharmacology for dynamic studies into animal behavior.

## Main

Photoactivatable or “caged” ligands are powerful tools for probing receptor signaling in living systems ^1–3^. Because photorelease is typically much faster than the biological processes under study, pre-equilibration of the caged molecule followed by photolysis can provide a robust stimulus that facilitates quantitative analysis of downstream processes. Although a few studies have achieved *in vivo* uncaging, none have demonstrated rapid responses that are desirable for studies into animal behavior ^4–8^.

The mu opioid receptor (MOR) is an inhibitory, G_i/o_-coupled G protein-coupled receptor (GPCR) that is widely expressed in the nervous system. MORs are the primary target of opioid analgesics such as morphine and fentanyl, and are activated by endogenous opioid neuropeptides enkephalin, beta-endorphin and dynorphin ^9,10^. How MOR activation in specific brain regions acutely impacts circuit activity and behavior, and how this changes with experience or adaptation to opioid drug use, remains unclear. To facilitate studies into opioid receptor signaling in the nervous system, we previously developed several caged opioid ligands, including analogues of the neuropeptide agonists [Leu^5^]-enkephalin (LE) and dynorphin A (1-8) (Dyn8), as well as the antagonist naloxone ^11–13^. Although LE activates MORs with nanomolar affinity, it also activates delta opioid receptors (DORs) in a similar concentration range ^9–11^. In addition, because peptides such as LE and, by extension, its caged variants, are rapidly degraded *in vivo* (t_1/2_ < 4 min) ^14^, currently available forms of caged LE are not readily compatible with *in vivo* experiments that investigate the role of MOR signaling in neural circuit function and animal behavior.

To enable *in vivo* experiments involving rapid activation of endogenous MORs, we developed a caged analogue of the MOR-selective peptide agonist [D-Ala^2^, N-MePhe^4^, Gly-ol]-enkephalin (DAMGO) ^15^. As DAMGO is an enkephalin derivative that was designed to resist proteolysis, it shares a cageable pharmacophore with LE and Dyn8. Our caged DAMGO design is based on our prior success in caging LE (**1**, **Figure 1**) by appending a caging group to the N-terminal tyrosine phenol to produce (carboxy-nitrobenzyl)-tyrosine-LE or CYLE (**2**) ^11^. Given the structural similarity between DAMGO (**3**) and LE, we reasoned that caging this site would similarly reduce the affinity of DAMGO for MOR. We chose to use the carboxy-nitroveratryl (CNV) caging group, as it includes a negatively charged carboxylic acid to facilitate solubility, and because it exhibits an absorbance tail that enables photolysis with 405 nm light. Importantly, we previously found that caging the *N*-Merminal tyrosine in LE with the CNV group (CNV-Y-LE) afforded rapid photorelease in brain slices to activate MORs with good sensitivity to UV light ^13^.

**Figure 1.**
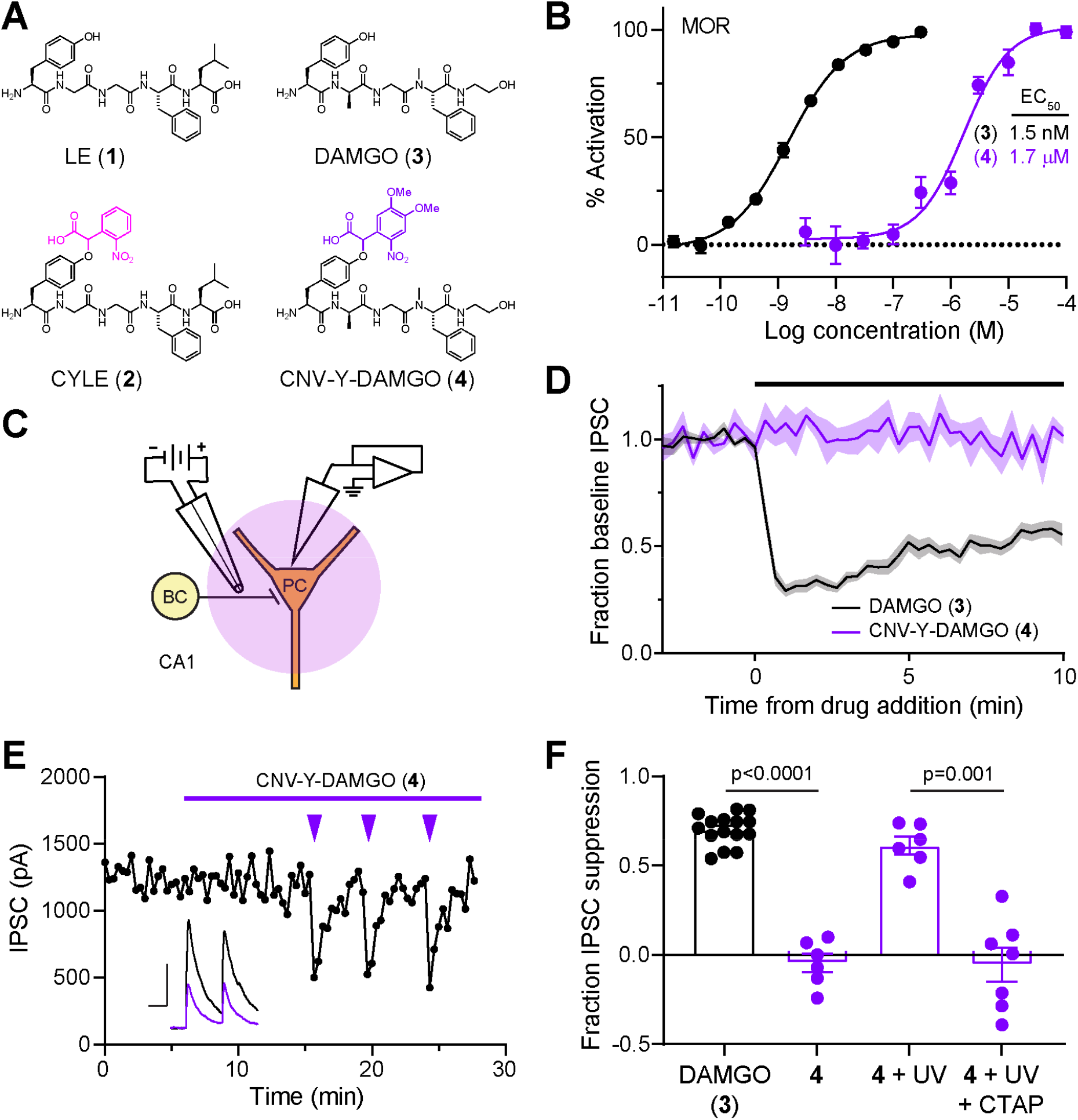
Design and validation of CNV-Y-DAMGO *in vitro* and *ex vivo*. (**A**) Design of a caged DAMGO based on caged [Leu^5^]-enkephalin. (**B**) Dose-response curves were obtained using a Glo-Sensor assay of cAMP signaling in HEK293T cells (n=12 wells per data point). Data were normalized to the maximal response to DAMGO (1 μM) and are expressed as the mean ± SEM. (**C**) Schematic of the experimental configuration for photouncaging of CNV-Y-DAMGO while recording electrically-evoked inhibitory synaptic transmission from CA1 hippocampal basket cells (BC) impinging on pyramidal cells (PC). (**D**) Baseline-normalized, average inhibitory post-synaptic current (IPSC) amplitude over time during bath application of DAMGO (1 μM, n = 15 cells from 14 mice) or CNV-Y-DAMGO (1 μM, n = 6 cells from 4 mice). (**E**) Representative example of IPSC amplitude over time during bath application of CNV-Y-DAMGO (1 μM) and repeated photolysis using 355 laser flashes (84 mW, 5 ms). **Inset:** Example IPSCs immediately before (black) and after (purple) CNV-Y-DAMGO uncaging. Scale bars: x = 25 ms, y = 500 pA. (**F**) Summary data comparing the fraction of baseline IPSC suppression in response to either DAMGO (1 μM) bath application, CNV-Y-DAMGO (1 μM) bath application, CNV-Y-DAMGO uncaging, and CNV-Y-DAMGO uncaging in the presence of the mu-selective antagonist CTAP (1 μM) (DAMGO (n = 15 cells from 14 mice), CNV-Y-DAMGO (n = 6 cells from 4 mice), CNV-Y-DAMGO + UV (n = 6 cells from 4 mice), CNV-Y-DAMGO + UV + CTAP (n = 7 cells from 2 mice)). p values were determined with a Mann-Whitney test.

Our three-step synthesis involves selective functional group modifications of commercially available DAMGO (**3**) to afford milligram quantities of CNV-Y-DAMGO (**4**) in ~45% overall yield (**Figure S1**). We evaluated the activity of CNV-Y-DAMGO at MORs using the GloSensor assay of G_α_ signaling in HEK293T cells (**Figure 1B**). Whereas we found the EC_50_ of DAMGO to be 1.5 nM in this assay, the EC_50_ of CNV-Y-DAMGO was 1.7 μM, which indicates that inactive concentrations of CNV-Y-DAMGO can be photoconverted to near saturating concentrations of DAMGO. We confirmed that CNV-Y-DAMGO cleanly releases DAMGO in aqueous solution by mass spectrometry, that it is completely stable in the dark for at least 24 h, and found that it undergoes photolysis at a rate ~2.25x that of MNI-glutamate, a caged glutamate derivative commonly used in brain slices (**Figure S2**).

To validate CNV-Y-DAMGO at native opioid receptors, we performed whole-cell voltage clamp recordings in mouse hippocampal brain slices, where electrically-evoked synaptic inhibition in the perisomatic region of CA1 pyramidal neurons is strongly suppressed by both MOR and DOR agonists ^13,16^ (**Figure 1C**). We first verified the inactivity of CNV-Y-DAMGO by recording inhibitory post-synaptic currents (IPSCs). Whereas bath application of DAMGO (1 μM) suppressed IPSC amplitude by ~70%, CNV-Y-DAMGO (1 μM) had no discernable effect on synaptic transmission (**Figure 1D**) (DAMGO: 0.70 ± 0.02; CNV-Y-DAMGO: −0.046 ± 0.05; p < 0.0001, Mann-Whitney test). We next found that 5 ms flashes of 355 nm light, applied 2 sec prior to an electrical stimulus, rapidly suppressed inhibitory synaptic transmission, and that this reversed over the course of several minutes (**Figure 1E**). The optically-evoked suppression of synaptic transmission was completely blocked by the selective MOR antagonist CTAP (**Figure 1F**) (CNV-Y-DAMGO + UV: 0.61 ± 0.05; CNV-Y-DAMGO + UV + CTAP: −0.05 ± 0.10, p = 0.001, Mann-Whitney test).

We pursued *in vivo* uncaging using optofluidic cannulas that deliver liquid and light to the same site, implanted above the left ventral tegmental area (VTA), where MOR agonists activate dopamine neurons through a disinhibitory mechanism ^17^ (**Figure 2A, B**). The resulting opioid-driven mesolimbic dopamine release leads to an increase in locomotion ^18^. Our experimental design is presented in **Figure 2C**. Based experiments with DAMGO that peaked 10-20 min post-infusion (**Figure S3**), we applied flashes of 375 nm light 15 min after infusion of CNV-Y-DAMGO (200 μM, 0.5 μl). For each mouse, we quantified both the instantaneous velocity (5 second bins), and the distance travelled in 15-minute windows before and after light application.

**Figure 2.**
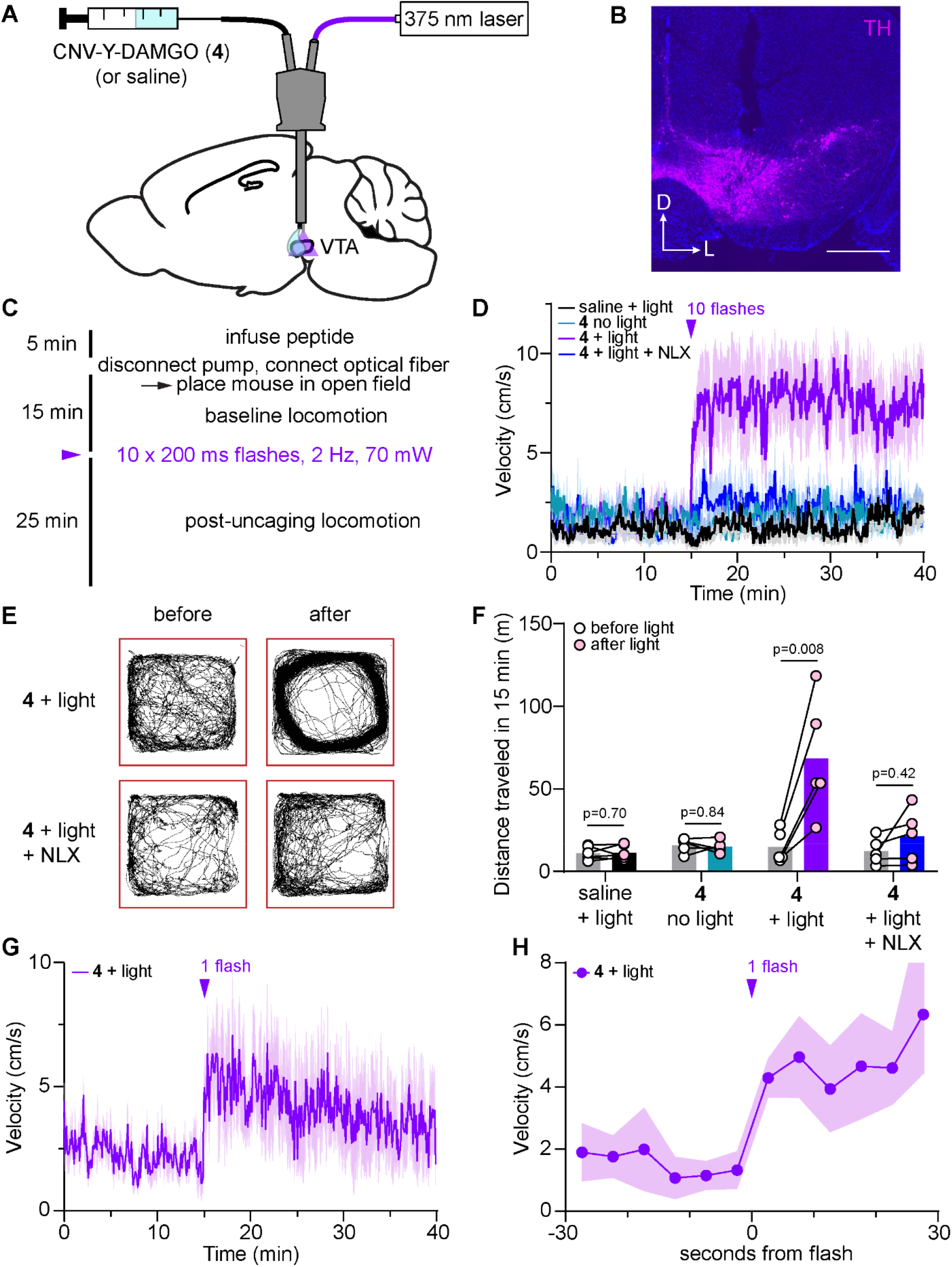
*In vivo* uncaging in the ventral tegmental area with CNV-Y-DAMGO. (**A**) Schematic of the experimental configuration for photo-uncaging through an optofluidic cannula. (**B**) Example image of cannula placement above the VTA, indicated by immunohistochemical labeling of tyrosine hydroxylase (TH). Scale bar = 500 μm. (**C**) Experimental timeline. (**D**) Average movement velocity vs. time for a cohort of mice treated with either saline and light (10 flashes, n=6 mice), CNV-Y-DAMGO without light (n = 5 mice), CNV-Y-DAMGO with light (n = 5 mice), and CNV-Y-DAMGO with light in the presence of the opioid antagonist naloxone (NLX) (n = 5 mice). The time of the uncaging stimulus is indicated by the purple arrowhead. (**E**) Example maps of open field locomotor activity before and after uncaging. (**F**) Summary data of distance traveled in the 15 minutes before and after application of the 10-flash uncaging stimulus (from the data shown in **E**). p values were determined with the Wilcoxon ranked sum test. (**G**) Average movement velocity vs. time upon exposure to a single light flash (n = 5 mice). (**H**) Same data as in **G**, but zoomed in around the uncaging stimulus.

Importantly, light alone had no effect on movement velocity (**Figure 2D**), and CNV-Y-DAMGO did not exhibit any residual activity at this dose when illumination was omitted. In contrast, pairing CNV-Y-DAMGO infusion with strong UV illumination (10 x 200 ms flashes, 1 Hz, 70 mW) produced a rapid increase in movement velocity that lasted over 25 min (**Figure 2D, E**). Systemic administration of the opioid antagonist naloxone (NLX, 3 mg/kg) strongly attenuated the locomotor response to DAMGO photorelease. These results are summarized in **Figure 2F**. A less intense optical stimulus (1 x 200 ms flash, 70 mW) produced a smaller, transient increase in locomotor behavior (**Figure 2G**). Strikingly, in both cases, locomotion was observed to increase within seconds of the flash, demonstrating that endogenous opioid receptors can rapidly modulate neural circuit function and behavior on this timescale (**Figure 2H**).

*In vivo* photopharmacology is currently an area of great research interest ^3,19–21^. A growing number of studies have used photoswitchable ligands to manipulate both genetically engineered and endogenous receptors *in vivo* with light ^22–28^. However, neither photoswitches nor caged ligands have been shown to trigger a near-instantaneous behavioral change such as that observed here ^4–8^. Behavior and physiology measurements typically require a several-minute delay for drugs to diffuse and equilibrate after local administration through cannulas. The ability to present a ligand to endogenous receptors with sub-second temporal precision is a key advantage of using photopharmacological tools *in vivo*, in addition to the spatial targeting afforded by light. Photoactivation can be paired with actions and events during behavioral experiments to study how receptor signaling in specific brain regions influences animal behavior and learning. Asking how this stimulus-response relationship is shaped by perturbing circuits and synapses (*e.g*. using DREADDs) can then be used to study the neural circuit mechanisms of drug action ^29^. In these ways, *in vivo* photopharmacology has the potential to revolutionize behavioral pharmacology research. Although the validation of CNV-Y-DAMGO for *in vivo* studies into opioidergic neuromodulation is an important step forward, this impending paradigm shift will require the development of an extensive, pharmacologically and optically diverse toolkit of *in vivo*-compatible photopharmacological probes, as well as specialized hardware optimized for delivery of both drug and light ^30–32^.

## Materials and Methods

### Chemical Synthesis and Characterization

High-resolution mass spectrometry data were obtained at the UCSD Chemistry and Biochemistry Mass Spectrometry Facility on an Agilent 6230 time-of flight mass spectrometer (TOFMS). CNV-Y-DAMGO was purified by reverse-phase chromatography to >99.9% purity, used as a mixture of diastereomers. DAMGO was obtained from HelloBio (HB2409) and TMSE-CNV-bromide (**6**) was synthesized according to a published procedure ^33^.

#### *N*-Boc-DAMGO (**5**)

To a stirred solution of DAMGO (**3**) (50 mg, 97 μmol) in 1,4-dioxane/water (3/1, 300 μL) in an amber glass vial at 22 °C, 1 M NaOH in water (97 μL, 97 μmol) was added followed by (Boc)2O (21 mg, 96 μmol) in 1,4-dioxane (50 μL). The reaction was monitored by HPLC. After 16 h, the reaction mixture was concentrated under vacuum and the residual was purified by C18 column chromatography (water/CH3CN, 95%/5% → 100%). Relevant fractions were combined and concentrated under vacuum to give *N*-Boc-DAMGO as a white solid (54 mg, 90%). LR-MS (ESI) *m/z* 636 [(M+Na)^+^, 100%]. HR-MS *m/z* 636.3005 [M+Na]^+^ (calcd for C31H43N5O8Na, 636.3004).

#### *N*-Boc-TMSE-CNV-Y-DAMGO

To an amber glass vial containing a stirred solution of *N*-Boc-DAMGO (**5**) (50 mg, 81 μmol) in anhydrous DMF (500 μL), NaH (60% in mineral oil, 3.3 mg, 82 μmol) was added slowly. The mixture was stirred at 0 °C for 0.5 h followed by the addition of TMSE-CNV-bromide (**6**, 34 mg, 81 μmol) in anhydrous DMF (100 μL) dropwise. The mixture was stirred under nitrogen and warmed up to 22 °C. The reaction was monitored by HPLC. After 3 h, a drop of MeOH was added to quench the reaction. The reaction mixture was concentrated under vacuum and the residual was purified by C18 column chromatography (water/CH3CN, 95%/5% → 100%). Relevant fractions were combined and concentrated under vacuum to give *N*-Boc-TMSE-CNV-DAMGO as a white solid (67 mg, 87%). LR-MS (ESI) *m/z* 975 [(M+Na)^+^, 100%]. HR-MS *m/z* 975.4145 [M+Na]^+^ (calcd for C_46_H_64_N_6_O_14_SiNa, 975.4142).

#### CNV-Y-DAMGO (**4**)

To an amber glass vial containing a stirred solution of *N*-Boc-TMSE-CNV-DAMGO (60 mg, 63 μmol) in DCM (200 μL), TFA (400 μL) was added. The reaction was monitored by HPLC. After 8 h, the reaction mixture was concentrated under vacuum and the residual was purified by C18 column chromatography (water/CH3CN, 95%/5% → 100%). Relevant fractions were combined and concentrated under vacuum to give CNV-Y-DAMGO as a white solid (27 mg, 58%). LR-MS (ESI) *m/z* 753 [(M+H)^+^, 60%], 775 [(M+Na)^+^, 100%]. HR-MS *m/z* 753.3086 [M+H]^+^ (calcd for C36H45N6O12, 753.3090).

### *In vitro* uncaging and dark stability

To determine dark stability, CNV-Y-DAMGO (**4**) (1 mM) was dissolved phosphate-buffered saline (PBS, pH 7.2) and left in the dark for 24 h. Comparison of samples taken at 0 and 24 h by HPLC-MS (1260 Affinity II, Agilent Technologies, Santa Clara, CA, USA) revealed no obvious decomposition or conversion to DAMGO. In addition to determining the chemical composition of the uncaging product by HPLC-MS, the initial photolysis rate of CNV-Y-DAMGO was compared to MNI-Glutamate using HPLC in response to illumination with a 375 nm laser. The concentrations of CNV-Y-DAMGO and MNI-glutamate were adjusted to match their optical densities at the photolysis wavelength of 375 nm. Solutions of CNV-Y-DAMGO (0.4 mM) and MNI-glutamate (0.5 mM) dissolved in PBS buffer (pH 7.2) were placed in 1 mL glass vials with stir bars and illuminated at a light intensity of 10 mW from a 375 nm laser (LBX-375-400-HPE-PPA, Oxxius, France) via an optical fiber (FT200UMT, 200 μm, 0.39 NA). The solutions were illuminated in 15 sec periods, after which samples were removed and analyzed by HPLC-MS using a linear gradient (water/MeCN 5%/95% → MeCN 100%, 0-8 min with 0.1% formic acid) and a C-18 column (4.6 × 50 mm, 1.8 μm) (Agilent). Each compound was assessed in triplicate. The integrals of the remaining caged molecule from each sample were normalized to the integral of the un-illuminated sample, averaged and plotted against time for the first two minutes of illumination. Linear regression provided a measurement of slope, and the ratio of the two slopes was used to determine the relative photolysis efficiency of CNV-Y-DAMGO in comparison to MNI-glutamate.

### GloSensor Assay

Human embryonic kidney 293T (HEK293T) cells were grown in complete DMEM (Dulbecco’s Modified Eagle’s Medium (DMEM, Invitrogen, Carlsbad, CA) containing 5% fetal bovine serum (Corning), 50 U/mL Penicillin-Streptomycin (Invitrogen), and 1 mM sodium pyruvate (Corning), and maintained at 37 °C in an atmosphere of 5% CO_2_ in 10 cm tissue culture (TC) dishes. When cell density reached ~70% confluence, growth medium was replaced with Opti-MEM (Invitrogen), followed by the addition of SSF-MOR plasmid ^34^, cAMP-dependent GloSensor reporter plasmid (−22F cAMP plasmid, Promega, E2301), and Lipofectamine 2000 (Invitrogen) in Opti-MEM. The dishes with transfection media were incubated at 37 °C in an atmosphere of 5% CO_2_ for 6 h before replacing the transfection medium with complete DMEM. After incubating at 37 °C in an atmosphere of 5% CO_2_ for 16 h, transfected cells were plated in poly-*D*-lysine coated 96-well plates at ~40,000 cells/well and incubated at 37 °C in an atmosphere of 5% CO_2_ for 16 h. On the day of assay, the medium in each well was replaced with 50 μL of assay buffer (20 mM HEPES, 1x HBSS, pH 7.2, 2 g/L *D*-glucose), followed by addition of 25 μL of 4x drug (DAMGO or CNV-Y-DAMGO) solutions for 15 min at 37 °C. Finally, 25 μL of 4 mM luciferin (Assay Reagent, Promega, E1291) supplemented with isoproterenol at a final concentration of 200 nM was added, and luminescence counting was performed after 25 min.

### Brain Slice Preparation

Animal handling protocols were approved by the UC San Diego Institutional Animal Care and Use Committee. Male and female postnatal day 15-35 mice on a C57Bl/6J background were anesthetized with isoflurane and decapitated. The brains were removed, blocked, and mounted in a VT1000S vibratome (Leica Instruments). Horizontal slices (300 μm) were prepared in ice-cold choline-ACSF containing (in mM) 25 NaHCO_3_, 1.25 NaH_2_PO_4_, 2.5 KCl, 7 MgCl_2_, 25 glucose, 0.5 CaCl_2_, 110 choline chloride, 11.6 ascorbic acid, and 3.1 pyruvic acid, equilibrated with 95% O_2_/5% CO_2_. Slices were transferred to a holding chamber containing oxygenated artificial cerebrospinal fluid (ACSF) containing (in mM) 127 NaCl, 2.5 KCl, 25 NaHCO_3_, 1.25 NaH_2_PO_4_, 2 CaCl_2_, 1 MgCl_2_, and 10 glucose, osmolarity 290. Slices were incubated at 32 °C for 30 min and then left at room temperature until recordings were performed.

### Electrophysiology

All recordings were performed within 5 h of slice cutting in a submerged slice chamber perfused with ACSF warmed to 32 °C and equilibrated with 95% O_2_/5% CO_2_. Whole-cell voltage clamp recordings were made with an Axopatch 700B amplifier (Axon Instruments). Data were sampled at 10 kHz, filtered at 3 kHz, and acquired using National Instruments acquisition boards and a custom version of ScanImage written in MATLAB (Mathworks) ^35^. Cells were rejected if holding currents were more negative than −200 pA or if the series resistance (<20 MΩ) changed during the experiment by more than 20%. Patch pipets (2.5-3.5 MΩ) were filled with an internal solution containing (in mM) 135 CsMeSO_3_, 10 HEPES, 1 EGTA, 3.3 QX-314 (Cl^-^ salt), 4 Mg-ATP, 0.3 Na-GTP, and 8 Na_2_ phosphocreatine (pH 7.3, 295 mOsm/kg). Cells were held at 0 mV to produce outward inhibitory currents. Excitatory transmission was blocked by the addition to the ACSF of NBQX (10 μM) and CPP (10 μM).To electrically evoke IPSCs, stimulating electrodes pulled from theta glass with ~5 μm tip diameters were placed at the border between stratum pyramidale and stratum oriens nearby the recorded cell (~50-150 μm) and two brief pulses (0.5 ms, 50-300 μA, 50 ms interval) were delivered every 20 s.

### *Ex vivo* UV Photolysis

Uncaging was carried out during brain slice electrophysiology recordings using 5 ms flashes of collimated full-field illumination with a 355 nm laser using a custom light path, as previously described ^11^. Light powers in the text correspond to measurements of a 10 mm diameter collimated beam at the back aperture of the objective. Beam size exiting the objective onto the sample through a 60x objective was 3900 μm^2^. Caged peptide was circulated in ACSF at the indicated concentrations at a flow rate of 4 ml/min using a peristaltic pump.

### *In vivo* UV Photolysis

Male and female postnatal day 40 mice on a C57Bl/6J background were implanted unilaterally with an Optical Fiber Multiple Fluid Injection Cannula (OmFC, Doric Lenses) just above the left ventral tegmental area at the following coordinates (in mm, from bregma): AP −3.33 mm, ML −0.5, DV −4.15) at a 10° M/L angle. The fluid guide cannula (485 μm OD) length was 4.5 mm, the flat tip (FLT) optical fiber (200 μm, 0.22 NA) length was 4.8 mm, the optical fiber receptacle was a 1.25 mm zirconia ferrule (ZF1.25), and the fluid injector (170 μm OD, 100 μm ID) length was 5.0 mm. Mice were allowed to recover for at least 5 days post-surgery and acclimated to the room and experimenter for several days prior to experimentation. Mice were brought to the room ~30 min prior to the start of each experiment. Using a syringe pump (WPI) connected to the fluid injector, CNV-Y-DAMGO (200 μM) was infused for 5 min at a flow rate of 0.1 μL/min. Mice were allowed to move freely for the duration of drug delivery. The fluid injector was allowed to remain in place for 1 min following infusion, after which the dummy fluid injector was reinserted. The optical fiber was coupled to a fiber optic cable (200 μm, 0.22 NA) connected to a 375 nm laser (Vortran Stradus), which controlled by an Arduino Uno using a custom code to control pulse duration and frequency. Mice were placed in a square enclosure (18 x 18 cm) where their position was video recorded at a 30 fps using a webcam (Logitech) fed into Smart 3.0 video tracking software (Panlab). Following a 15 min baseline period, the indicated number of light flashes (200 ms, 1 Hz) were applied at the indicated powers, after which mice were tracked for an additional 25 min. Reported light power refers to measurements from the ferrule tip of the laser-coupled optical fiber (200 μm, 0.22 NA) that connects to the implanted fiber. In some experiments CNV-Y-DAMGO was replaced with saline only, the light flashes were omitted, or naloxone (3 mg/kg, intraperitoneal) was administered immediately before beginning the infusion.

### Histology

Mice were anesthetized with isoflurane via the open drop method and perfused with 20 mL of PBS, followed by 20 ml of chilled 4% PFA in 1x PBS. The brains were collected, placed in 4% PFA overnight at 4°C, and transferred to a 30% sucrose solution for at least 18 h prior to sectioning. Brains were sliced into 40 μm sections using a sliding microtome (Thermo Scientific Microm HM 450) and collected in PBS in 24-well plates. Sections selected for immunochemistry were blocked in PBS-Triton (0.3% TritonX-100 in 1x PBS) + 5% donkey serum for 2 h at room temperature. The sections were then incubated with rabbit anti-tyrosine hydroxylase (Millipore, AB152; 1:000) in PBS-Triton + 1% donkey serum at 4°C overnight while shaking. Following primary incubation, sections were washed 5x at 10 min per wash in PBS, then incubated with Alexa Fluor 647 donkey anti-rabbit IgG (1:1000) in PBS-Triton + 1% donkey serum for 2 h at room temperature for secondary staining. Following secondary incubation, sections were washed 5x at 10 min each in PBS, then mounted on glass slides using Vectashield Antifade Mounting Medium with DAPI (H-1200). Images of the slices were acquired on a Keyence (BZ-X710 series) at 10x magnification.

### Data Analysis

Electrophysiology data were analyzed in Igor Pro (Wavemetrics). Peak current amplitudes were calculated by averaging over a 2 ms window around the peak of the IPSC. To determine magnitude of modulation by DAMGO uncaging (%IPSC suppression), the IPSC peak amplitude immediately after a flash was divided by the average peak amplitude of the three IPSCs preceding the light flash. The effects of drugs on IPSC suppression were calculated as the average %IPSC suppression 2-3 min after drug addition. Video measurements of mouse position over time were analyzed in either 5 or 1 s bins for instantaneous velocity determination, and in 15 min bins before and after uncaging for calculating total distance traveled (Smart 3.0 video tracking software, Panlab). Summary values are reported as mean ± SEM. All data are treated as nonparametric and specific statistical tests and corrections are described for each figure in the text and figure legends.

## Acknowledgements

We would like to thank Eric Berg for technical support, Jeffry Isaacson for helpful discussions, and Janie Chang-Weinberg for manuscript editing.

## Funding

This work was supported by the National Institute of Neurological Disorders and Stroke and the National Institute of Mental Health (U01NS113295 to M.R.B.), the National Institute of General Medical Sciences (T32GM007240 to X.J.H.), the Brain & Behavior Research Foundation (M.R.B), the Esther A. & Joseph Klingenstein Fund & Simons Foundation (M.R.B), and the Rita Allen Foundation (M.R.B.).

## Figures and Figure Legends

**Figure S1.**
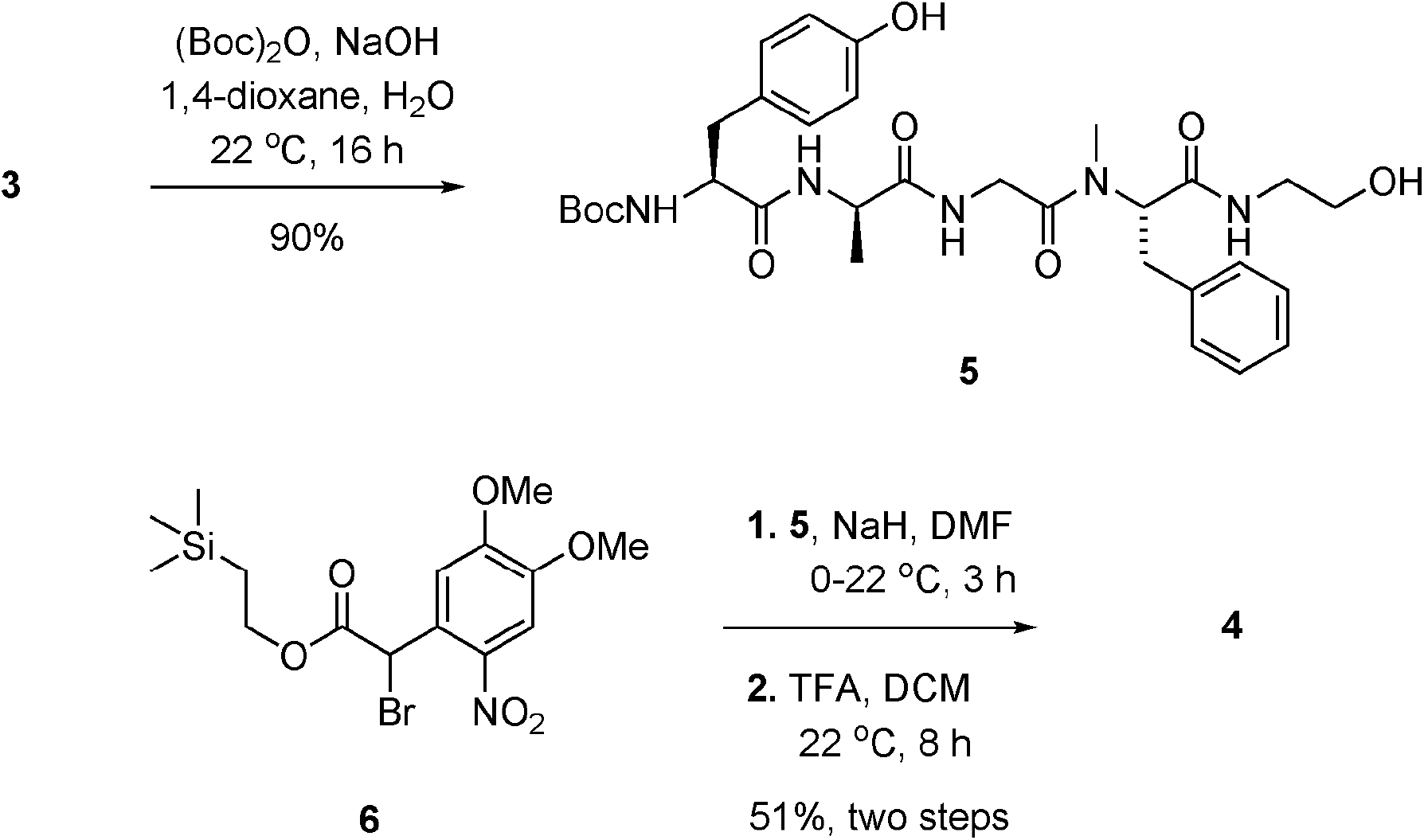
Synthesis of CNV-Y-DAMGO (4).

**Figure S2.**
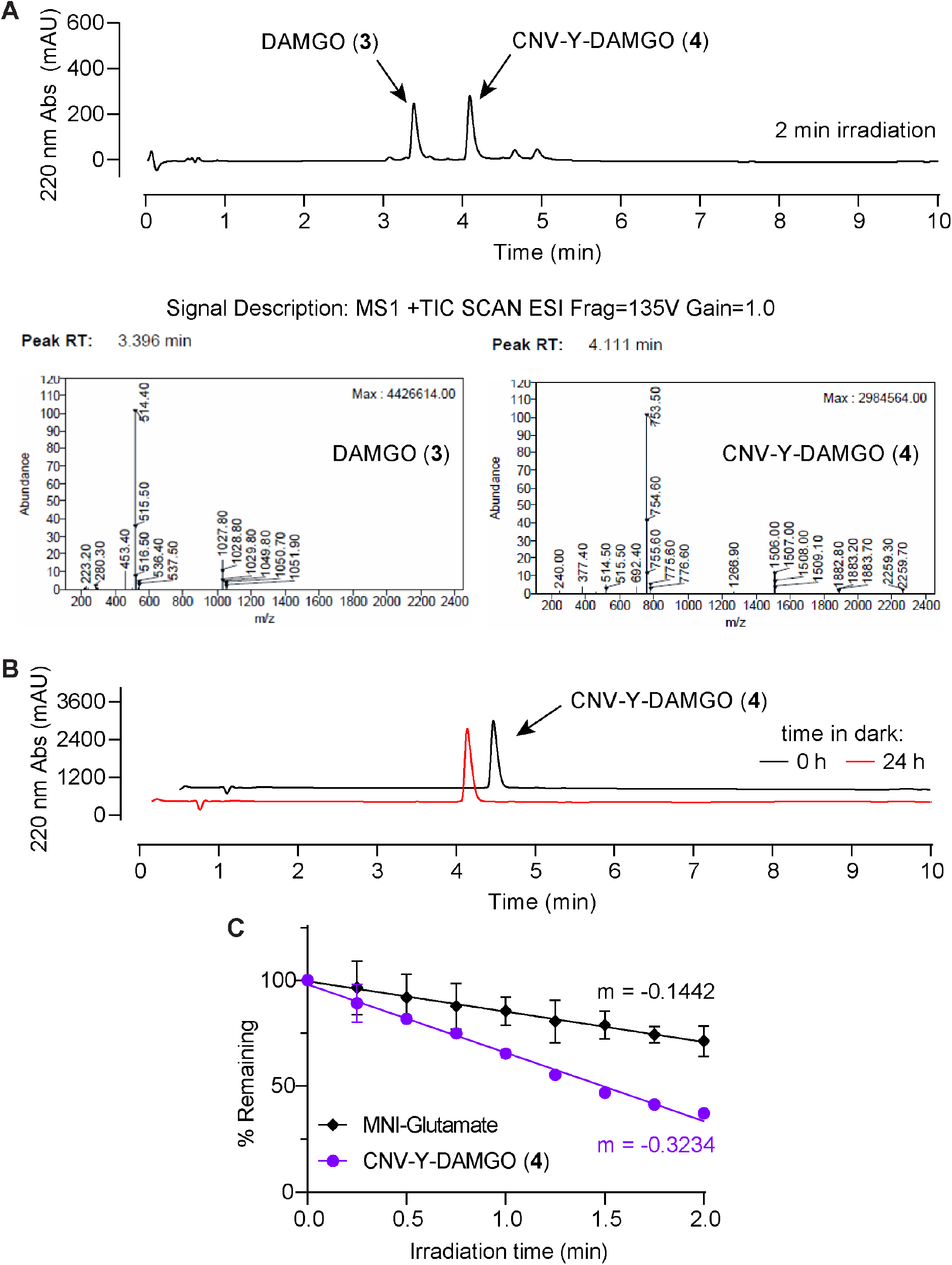
*In vitro* characterization of CNV-Y-DAMGO (4) photolysis and dark stability. (**A**) Waterfall plot of HPLC chromatograms demonstrating complete photoconversion of CNV-Y-DAMGO to DAMGO upon illumination with 375 nm light in PBS, pH 7.2. (**B**) Enlarged HPLC chromatogram after 2 min of illumination showing that photolysis of CNV-Y-DAMGO in PBS with 375 nm light releases a single major product (top) that is confirmed to be DAMGO by mass spectroscopy (bottom). (**C**) HPLC chromatograms showing that a solution of CNV-Y-DAMGO in PBS is stable for at least 24 hours in the dark. (**D**) Summary of MNI-Glutamate and CNV-Y-DAMGO photouncaging reactions over time, as measured by HPLC (n=3 samples per condition). Samples were optical density-matched at 375 nm in PBS and illuminated with 375 nm laser irradiation.

**Figure S3.**
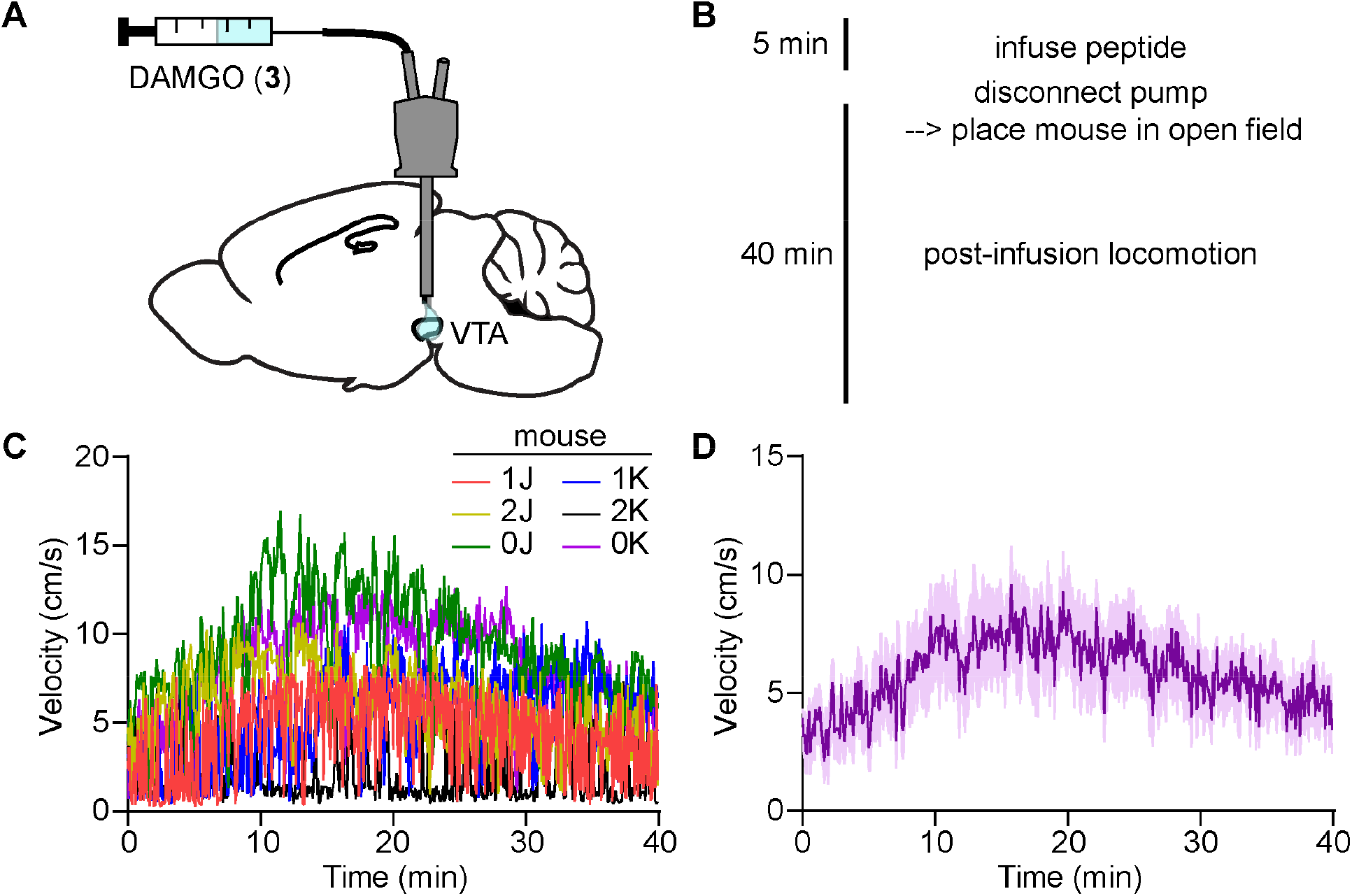
Identification of the time-course of DAMGO-driven locomotor behavior when infused unilaterally into the VTA. (**A**) Schematic of the experimental configuration for DAMGO infusion through an optofluidic cannula. (**B**) Experimental timeline. (**C**) Movement velocity vs. time in the open field for a cohort of mice after DAMGO (200 μM, 0.5 uL) infusion. (**D**) Average data from **C** (n=6 mice).

## Notes

**Competing Interests Statement** The authors declare no competing financial interests.

### Competing Interest Statement

The authors have declared no competing interest.

### Summary of Updates

In vivo uncaging data were added to the manuscript, in addition to additional authors. The title has changed to reflect the increased scope.

